# Ocular Response Functions reveal how ocular processes relate to neural activity

**DOI:** 10.1101/2024.11.19.624356

**Authors:** Quirin Gehmacher, Juliane Schubert, Aaron Kaltenmaier, Nathan Weisz, Clare Press

**Affiliations:** Department of Experimental Psychology, University College London, WC1H 0AP, United Kingdom; Department of Imaging Neuroscience, UCL Queen Square Institute of Neurology, University College London, WC1N 3AR, United Kingdom; Paris-Lodron-University of Salzburg, Department of Psychology, Centre for Cognitive Neuroscience, Salzburg, Austria; Neuroscience Institute, Christian Doppler University Hospital, Paracelsus Medical University, Salzburg, Austria

## Abstract

Oculomotor activity provides critical insights into cognition and health, with growing evidence demonstrating its involvement in various cognitive functions such as attention, memory, and sensory processing, and that it is a significant indicator of psychopathologies and neurological disorders. Despite its crucial importance across domains, the neural mechanisms supporting oculomotion have been underexplored, largely because eye movements are typically treated as artefacts to be removed from the neural signal. While useful for data cleaning, this approach risks discarding valuable information about oculomotor control, and there has been recent interest in modelling these signals instead to understand them. Using time-resolved regression methods with magnetoencephalography (MEG) and eye tracking during the resting state, we thus here sought to model ‘Ocular Response Functions’ (ORFs), that characterise the neural signatures of distinct oculomotor events, specifically saccades, blinks, and pupil dilation. We demonstrate the relationships between ocular action and neural activity (encoding), revealing a range of sensory cortical, cerebellar and frontal signatures preceding and following such ocular events. We conversely show how we can reconstruct ocular events from brain activity, and further apply resting-state derived ORFs to a passive listening task - demonstrating how some neural markers interpreted as directly related to sensation may in principle indirectly result from oculomotor contributions to task-related neural processing. By providing an accessible framework for examining the interplay between eye movements and neural processes, we offer a range of insights with potential applications across cognitive and clinical neuroscience.

## Introduction

Oculomotor activity offers a unique opportunity for understanding key aspects of cognition and health ^1^. A growing body of work demonstrates that eye movements are tightly coupled to a wide range of cognitive functions, including attention, memory, and visual and auditory processing ^2–17^ - serving both as a sensitive readout of ongoing cognitive states and, in some domains, as an active contributor to cognitive functions. Relatedly, oculomotor activity is increasingly recognised as an important indicator of psychopathology and neurological disorders, with links to conditions such as schizophrenia ^18–23^, dementia ^24–28^, and depression ^29–32^, and even phantom perceptions like tinnitus ^33–35^.

Despite this growing recognition, examining how oculomotor processes are related to neural activity has remained difficult. Conventionally, such ocular activity is removed from neural data - excluded or corrected through methods like Independent Component Analysis (ICA) - yet regression and system-identification methods can instead be used to model the neural response to ocular events directly, and in principle study rather than correct for them ^36–44^. For example, powerful toolboxes have been developed to simultaneously model ocular, alongside task (e.g. stimulus), events and thus methodologically start to disentangle these signals ^40^. However, disentangling that influence is still difficult: ocular events are mutually correlated - saccades, blinks, and pupil changes co-occur - and are and are themselves modulated by the task, each generating its own neural response ^45^. Time-resolved deconvolution can separate overlapping responses only where their relative timing varies ^40^ yet task-driven eye movements share their timing with the stimulus response, so the ocular and cognitive contributions become correspondingly hard to separate within the task itself. The ocular contribution is thus described, but not itself made available as an oculomotor-related neural signal that can be carried into a separate task and analysed alongside the cognitive processes of interest. As a result, we still need to understand better the neural processes generating and resulting from ocular action, as well as the extent to which the neural activity we attribute to a cognitive process reflects the concurrent activity of the eyes.

Here we introduce Ocular Response Functions (ORFs) to illuminate these interactions. Building on established time-resolved regression methods ^41–43,46^, we estimate, for each ocular feature, the neural impulse response that characterises its signature across the brain, deriving these functions from task-free resting-state data - that is, in the absence of stimulus-evoked activity that would otherwise couple ocular and cognitive responses in time - modelling saccades, blinks, and pupil dilation within a single design so that each function reflects the unique contribution of its event rather than a mixture of these correlated signals. Estimated in magnetoencephalography (MEG) source space, these functions resolve not only the time course but also the anatomical distribution of each ocular event’s neural signature. Convolving the ocular signals recorded during a separate task with these transfer functions then re-expresses ocular activity as a predicted neural signal, placing the ocular contribution on the same dimension as the brain recording. This makes it possible to test whether, and to what extent, oculomotor activity contributes to neural responses in tasks where eye movements are not ordinarily considered. The same functions can also be applied in reverse, to reconstruct ocular events from neural activity. This is hardest for sparse, binary events such as saccades and blinks: standard backward decoding is designed for continuous targets, and does not separate the contributions of correlated features very well ^47^, so their reconstruction has rarely been attempted. To recover such discrete events, we time-reverse each jointly-estimated response function and convolve it with the neural signal - a matched-filter operation that inherits the feature separation already built into the forward model. Thresholding the result then detects events directly, adapting the logic of spike detection from electrophysiology.

We present our findings in two stages. First, from simultaneous MEG and eye-tracking recorded during the resting state, we derive ORFs and characterise the distinct spatiotemporal signatures of saccades, blinks, and pupil dilation - both from data cleaned of ocular artefacts (ICA) and from uncorrected data - confirming that they support encoding (predicting neural activity from ocular events) and reconstruction (recovering ocular events from neural activity). We then apply the resting-state ORFs to a separate passive-listening task, in which participants heard tone sequences that differed in their statistical regularity. Re-expressing the task’s eye movements as a predicted neural signal, we find that some of the neural signature of auditory regularity tracking is likely a function of (small) eye movements. We present this as a proof of principle: a demonstration that oculomotor activity, expressed in neural space, carries information about ongoing cognition, and that its contribution to task-related neural activity can be examined directly. By providing a framework for examining the interplay between eye movements and neural processes, we hope our approach opens further avenues for both research and clinical applications.

## Results

In the following, we present the ‘Ocular Response Function’ (ORF) and organise the results into three sections. First, we outline the general steps for estimating the ORFs, providing a schematic overview of the approach, and how they can be used to predict brain activity from ocular action (encoding) or ocular action from brain data (reconstruction). Second, we show that ORFs account for variance across distinct brain regions, each with its own temporal profile, in data both with and without artefact correction (ICA and no-ICA). By convolving resting-state data with the corresponding ORFs, we demonstrate that brain activity can be predicted from ocular activity, and ocular activity from brain activity. Third, we apply the ORF approach to a passive-listening task, asking whether - and how - oculomotor processes contribute to neural activity in a setting where eye movements are not ordinarily considered.

### The ‘Ocular Response Function’

To establish a time-continuous and spatially resolved relationship between ocular features and neural activity, we first projected the continuous resting-state MEG data into anatomical source space using linearly constrained minimum variance (LCMV) beamformers. Then, for each source-space voxel, we fitted a forward encoding model using a deconvolution algorithm based on ^46^ (see Fig. 1A) as implemented in the Eelbrain toolkit ^41^. This model estimated the unique relationship between each of our three ocular features (pupil dilation, blinks, and saccades) and the neural activity at that specific brain location (see Fig. 1B). Fitting these models to task-free resting-state data yields eye-brain response functions, or ORFs, estimated in the absence of stimulus-evoked activity (see Fig. 1C). This process yields a unique ORF for each ocular feature at each voxel, reflecting the strength of the relationship at different time lags. For example, a value at +100 ms reflects the brain’s response following an ocular event, whereas a value at −100 ms reflects neural activity preceding that same event.

**Fig. 1:**
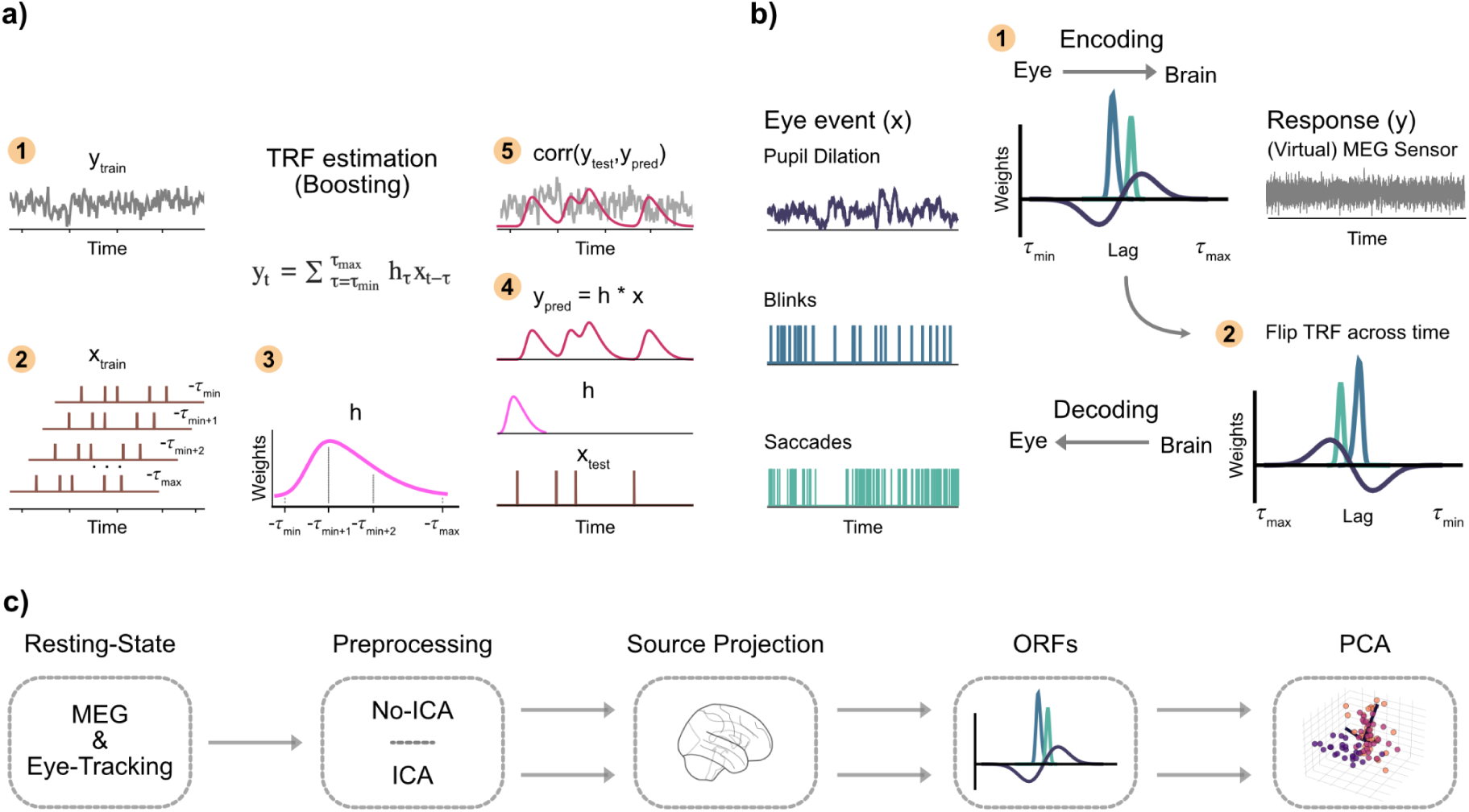
The Ocular Response Function. **a)** Eye (x) - response (y) relationships are estimated using deconvolution (Boosting), where neural activity (y_train_) is linked to an ocular action (x_train_, e.g. saccades) with time-shifted versions to capture the relationship across time with the neural signal. The TRF (filter kernel, h) represents how the neural signal is related to the eye event over time. This filter is then applied (using convolution) to new eye data (x_test_) to predict the neural response (y_pred_). Finally, the predicted response is compared to the actual neural activity (y_test_) to assess the model fit and evaluate how accurately the TRF captures the brain’s response dynamics. This TRF approach has been previously developed to establish the relationship between experimental events (x) and neural activity (y), but we replace the x vectors with those reflecting ocular action to generate Ocular Response Functions (ORFs). **b)** Specifically, ORFs leverage this approach by predicting resting-state brain activity from three key ocular features (pupil dilation, blinks, and saccades) using a forward encoding model. One ORF is generated per ocular feature per source-space location (virtual sensor). To reconstruct (decode) eye movements from brain data, the ORF is reversed along the temporal axis. **c)** Models were fitted to data derived from resting-state MEG recordings that were either cleaned of oculomuscular and cardiac artefacts (ICA) or had these components retained (no-ICA). In the present study, the continuous MEG data was first projected into source space. The ORFs were then estimated directly on the time series of each source-space voxel, resulting in anatomically localised transfer functions. Finally, principal component analysis (PCA) was applied to the resulting source-space ORFs to summarise the data and identify the dominant spatiotemporal patterns of eye-brain coupling.

Once the ORFs are estimated on training data, their performance is evaluated on held-out test data using a cross-validation procedure. For reconstruction, the same functions are reversed in time and applied to the neural data to predict ocular events, supporting a bidirectional account of eye-brain interactions. Models were fitted to source-projected data derived from MEG recordings either with or without prior ICA-based artefact correction, allowing us to differentiate the contributions of ocular artefacts from oculomotor-related neural activity in M/EEG recordings. For evaluation and application to other experimental tasks, we first convolve each ocular feature with the corresponding source-projected ORF - in a process analogous to the hemodynamic response function (HRF) convolution used in fMRI analysis - to generate a time course of predicted brain activity for each voxel, based on the respective eye-tracking data. We then correlate these predicted time courses with the actual source-projected brain recordings to assess encoding performance (see Fig. 2A). Conversely, by convolving the source-projected brain activity with the time-flipped ORFs, we generate a predicted time course of each ocular feature, which is then correlated with the actual eye-tracking recordings to assess reconstruction performance (see Fig. 2B).

**Fig. 2:**
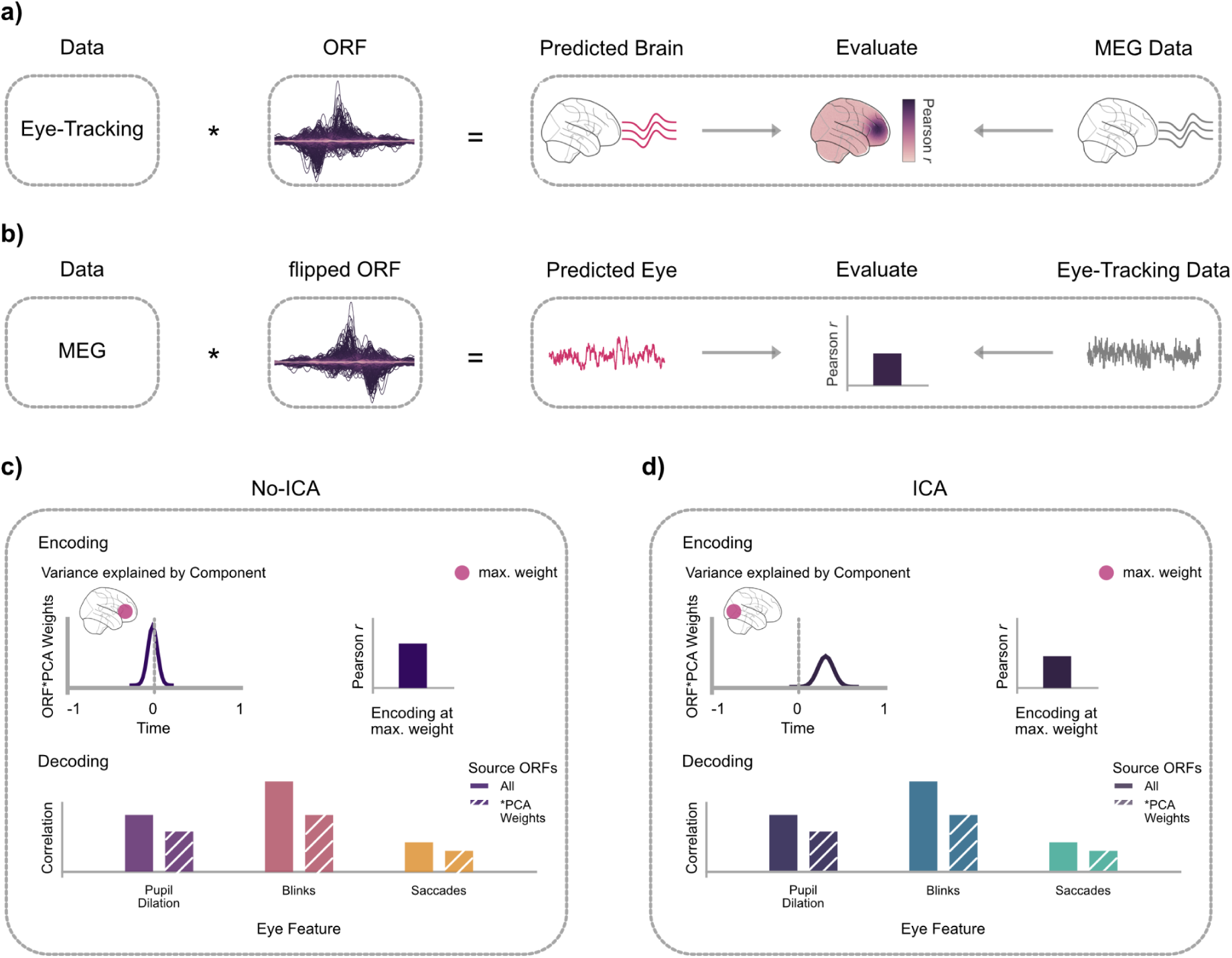
The approach for obtaining encoding and reconstruction results. **a)** Encoding evaluation. A predicted neural time course is generated by convolving ocular data with a derived ORF. This prediction is then correlated with the actual neural recordings to assess model fit at each brain location. **b)** Reconstruction evaluation. A predicted ocular time course is generated by convolving neural data with a time-flipped ORF. This prediction is then correlated with the actual ocular recordings to assess reconstruction performance. **c & d)** Schematic of the PCA-based analysis and the patterns it is expected to reveal. The diagrams illustrate how PCA is used to identify dominant components from the source-space ORFs, allowing for a comparison between data without (c) and with (d) ocular artifact correction. The illustration reflects the hypothesis that uncorrected data would be dominated by large-scale frontal artifacts originating from eye movements, while corrected data would reveal more distributed neural sources.

To summarise these high-dimensional results and identify the dominant patterns of eye-brain coupling, we applied principal component analysis (PCA) to each participant’s source-projected ORFs. The resulting spatial weight maps for the principal components were then averaged across all participants. For our quantitative analysis, we identified the location of maximum weight within these group-averaged maps. This group-defined location was then used to extract the cross-validated encoding effect from each individual participant’s data. In the following, we present distinct ORF time courses for the first component, along with encoding effects at the locations of maximum weight. Additionally, reconstruction effects are demonstrated by predicting eye movement patterns using all source-projected ORFs, either collectively or weighted by the PCA matrices of the first component (results for all other component-based models can be found in Supplementary Material, Supplementary Figures, Fig. S3).

### Ocular response functions: encoding and reconstruction during the resting state

First, we established ORFs by modeling resting-state brain activity from three ocular features: pupil dilation, blinks, and saccades. To visualise the dominant spatiotemporal patterns in these relationships, we used PCA, applying it to the source-space ORFs from data with and without ocular artifact correction (ICA and no-ICA, respectively).

In brain data without ICA preprocessing, the first principal component (PC1) for each ocular feature was generally characterised by strong loadings over frontocentral or early sensory regions. After applying ICA, these loadings typically maintained the early sensory responses, alongside now clearer contributions localised to the cerebellum, revealing patterns that were likely previously obscured by artifacts from eyeball rotation. Below, we present the temporal and spatial patterns for PC1 for both preprocessing strategies.

#### No-ICA Preprocessing

For pupil dilation, the first component (explained variance: *μ* = 31.7%, sd = 8.1%) was characterised by primary loadings over posterior medial regions, including the precuneus. The associated time course was dominated by a large peak at approximately −400 ms, suggesting that activity in these regions precedes changes in pupil diameter (encoding at the group-defined peak voxel of this component, *r* = 0.03, *t*(28) = 6.26, *p* < .001, *d* = 1.16; see Fig. 3A). For eye blinks, PC1 (explained variance: *μ* = 61.1%, *sd* = 20.2%) showed maximal loadings in frontocentral regions, consistent with ocular artifacts, and peaked near-instantaneously around 0 ms (encoding at the component’s peak, *r* = 0.35, *t*(28) = 13.22, *p* < .001, *d* = 2.45; see Fig. 3B). Finally, for saccades, PC1 (explained variance: *μ* = 42.6%, sd = 20.2%) had widespread loadings with a maximum in occipital visual areas, including the calcarine sulcus. Its time course showed a sharp peak at approximately +100 ms, indicating a strong visual response immediately following saccade execution (encoding at the peak voxel, *r* = 0.08, *t*(28) = 7.69, *p* < .001, *d* = 1.43; see Fig. 3C).

**Fig. 3:**
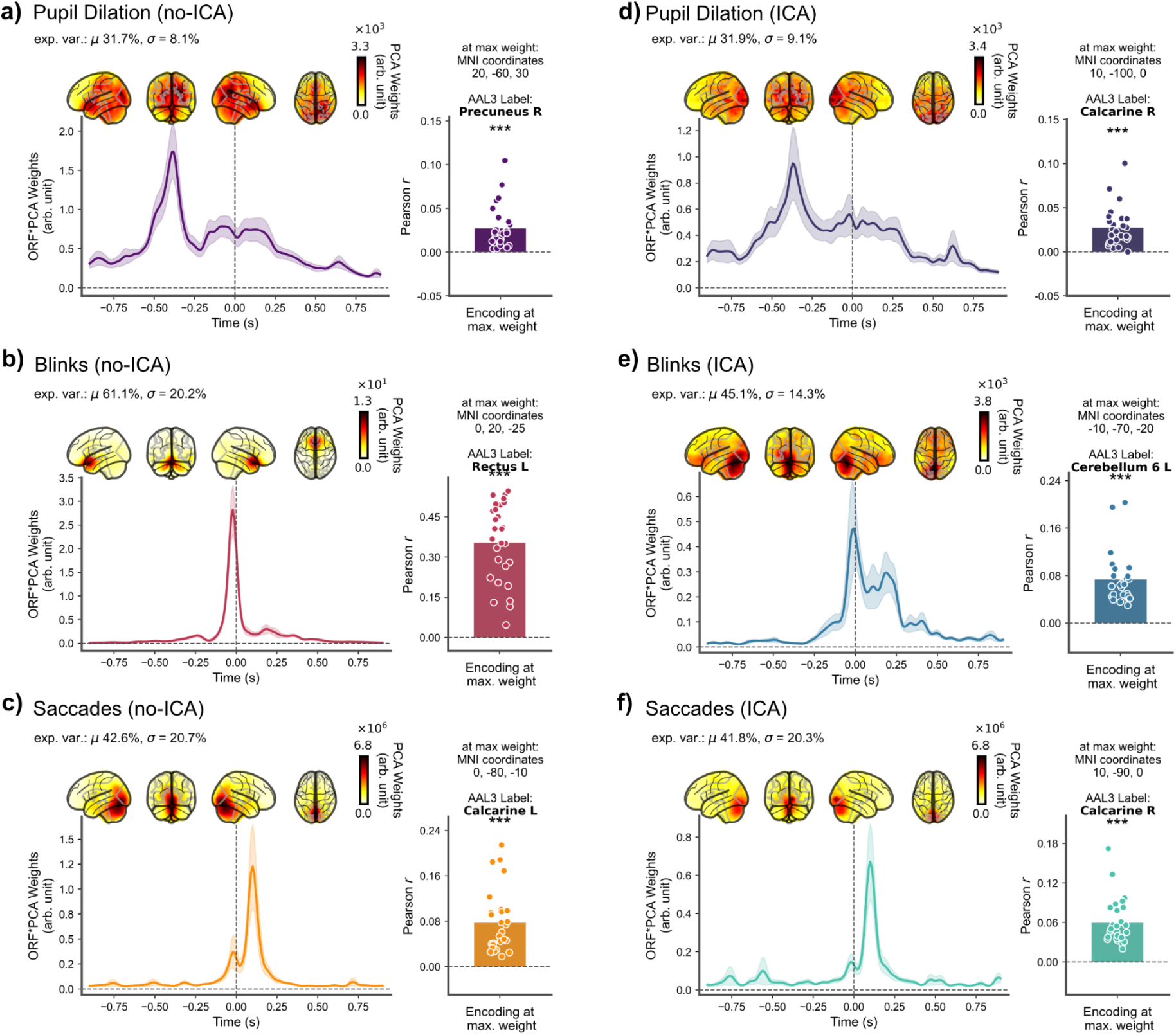
Encoding results for the first principal component (PC1) of the Ocular Response Functions. Results are shown for data without (no-ICA) and with (ICA) ocular artifact correction. Without ICA, activity is dominated by frontocentral, posterior medial, and occipital activity. After ICA, topographies shift to retain occipital loadings and additionally reveal clearer neural sources in cerebellar regions. **a-c)** No-ICA Results: **a)** Pupil dilation: The component showed primary loadings over posterior medial regions (e.g., precuneus) and peaked at ∼ −400 ms. **b)** Eye blinks: The component was localised to frontocentral regions, consistent with ocular artifact, and peaked around 0 ms. **c)** Saccades: The component loaded maximally on the occipital cortex (e.g., calcarine sulcus) with a peak at ∼ +100 ms, reflecting a strong visual response. **d-f)** ICA-Cleaned Results: **d)** Pupil dilation: After artifact correction, the component’s primary loading shifted to occipital visual areas, with a peak at ∼ −300 ms. **e)** Eye blinks: The component was now localised to the cerebellum and exhibited a more complex response with a secondary peak around 150-200 ms. **f)** Saccades: The component remained localised to the occipital visual cortex with a peak at ∼ +100 ms, confirming a robust, non-artifactual neural source. *General Notes*: Shaded error bars represent 95% CI’s. Anatomical labels were obtained using the automated anatomical labelling atlas 3 (AAL3). Statistics were performed using one-sample Student’s t-tests. ‘n.s.’: not significant; ‘\**’*: *p* < 0.05; *‘*\*\*’: *p* < 0.01; ‘***’: *p* < 0.001. *N* = 29.

#### ICA Preprocessing

After ICA correction, the spatial topographies no longer reflected any frontocentral loadings, but retained the sensory sources which were accompanied now by localisations to the cerebellum. For pupil dilation, PC1 (explained variance: *μ* = 31.9%, *sd* = 9.1%) now maximally loaded over occipital visual areas. The time course showed a prominent peak around −300 ms, suggesting that activity in the visual cortex precedes upcoming changes in pupil size (encoding effect, *r* = 0.03, *t*(28) = 6.77, *p* < .001, *d* = 1.26; see Fig. 3D). For eye blinks, PC1 (explained variance: *μ* = 45.1%, sd = 14.3%) was now localised to the cerebellum. The time course exhibited an initial near-instantaneous peak, but now also showed a more pronounced second activation around 150-200 ms, suggesting a complex, delayed response to blinks after ocular artifacts were suppressed (encoding effect: *r* = 0.07, *t*(28) = 9.45, *p* < .001, *d* = 1.76; see Fig. 3E). For saccades, PC1 (explained variance: *μ* = 41.4%, sd = 20.3%) showed a similar pattern to the no-ICA data, with primary loadings in occipital visual areas and a response peak around +100 ms - showing saccades are followed abruptly by changes in visual responses. The stability of this occipital component post-ICA suggests it reflects a robust neural response to saccadic reafference, clearly separable from frontal muscle artifacts (encoding effect: *r* = 0.06, *t*(28) = 9.47, *p* < .001, *d* = 1.76; see Fig. 3F).

The spatial and temporal patterns for the second and third principal components generally showed loadings over parietal and sensory regions. For detailed results and visualisations for these components, please refer to the Supplementary Material (Supplementary Figures, Fig. S1&2).

Next, we assessed how well each ocular feature could be reconstructed from the resting-state neural data. These predictions were generated by convolving the source-projected brain data with the corresponding time-flipped ORFs. We tested this using either the full source-space data (’All’) or data restricted to the patterns of the top three principal components (’PC1’, ‘PC2’, ‘PC3’), as well as their combination (’PCA’). Reconstruction performance was quantified by correlating the predicted and actual eye-tracking time series, using Pearson’s *r* for continuous data (pupil dilation) and the Matthews Correlation Coefficient (*MCC*) for binary events (blinks and saccades). In the main text, we focus on the results from the ‘All’ source model, which represents the total reconstructed information, and the ‘PC1’ model, which reflects information contained in the single most dominant spatiotemporal component.

As detailed in Figure 4, all three ocular features were reconstructed at levels convincingly above chance (t values ranging from 8 to 21) in both the no-ICA and ICA-cleaned data. In all cases, performance was highest when using the full source model (’All’) and, while reduced, remained highly significant when using only the first principal component (’PC1’).

**Fig. 4:**
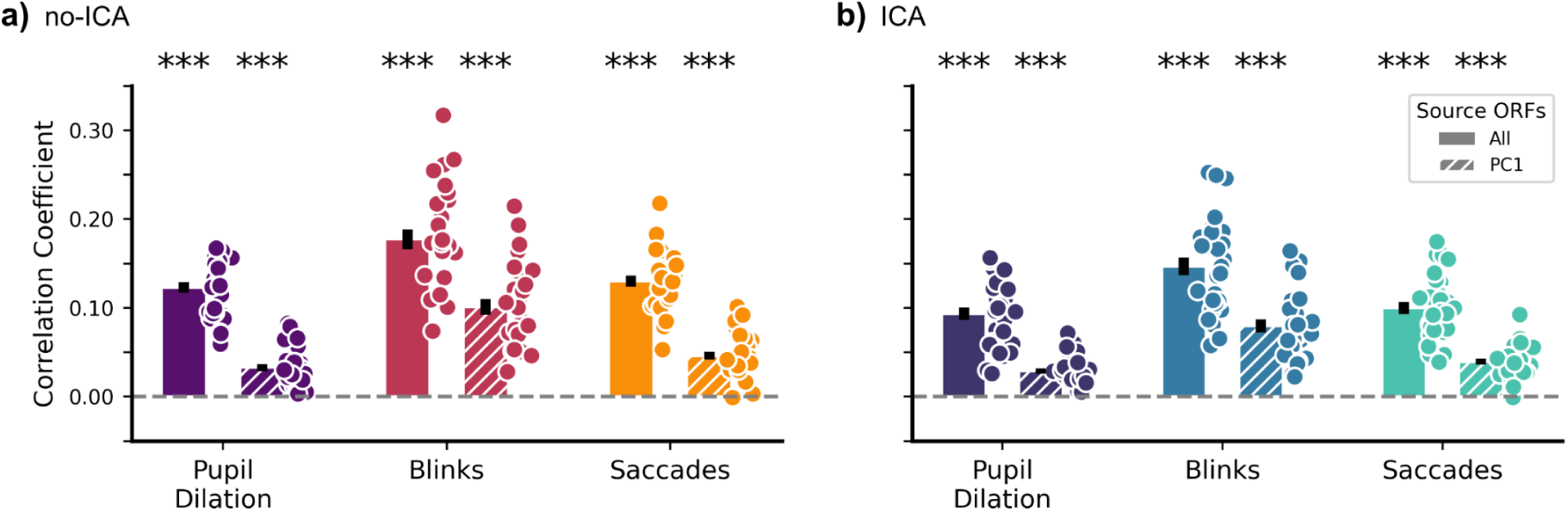
Reconstruction performance for ocular features from resting-state data. Reconstruction was performed using all source-space ORFs (’All’) or only those weighted by the first principal component (’PC1’). **a)** No-ICA preprocessing: Using ‘All’ sources yielded significant decoding for all features (Pupil: *r* = 0.12, *t*(28) = 20.73, *p* < .001, *d* = 3.85; Blinks: *MCC* = 0.18, *t*(28) = 15.91, *p* < .001, *d* = 2.95; Saccades: *MCC* = 0.13, *t*(28) = 21.37, *p* < .001, *d* = 3.97). Using only ‘PC1’ resulted in lower but still highly significant performance (Pupil: *r* = 0.03, *t*(28) = 8.66, *p* < .001, *d* = 1.61; Blinks: *MCC* = 0.10, *t*(28) = 11.62, *p* < .001, *d* = 2.16; Saccades: *MCC* = 0.05, *t*(28) = 10.25, *p* < .001, *d* = 1.90). **b)** ICA preprocessing: After ICA, decoding with ‘All’ sources remained strong (Pupil: *r* = 0.09, *t*(28) = 13.01, *p* < .001, *d* = 2.42; Blinks: *MCC* = 0.15, *t*(28) = 14.92, *p* < .001, *d* = 2.77; Saccades: *MCC* = 0.10, *t*(28) = 15.05, *p* < .001, *d* = 2.79). Performance with ‘PC1’ was also significant (Pupil: *r* = 0.03, *t*(28) = 9.31, *p* < .001, *d* = 1.73; Blinks: *MCC* = 0.08, *t*(28) = 10.69, *p* < .001, *d* = 1.99; Saccades: *MCC* = 0.04, *t*(28) = 11.32, *p* < .001, *d* = 2.10). Statistics reflect one-sample Student’s t-tests against zero. Error bars reflect standard error of the mean. ‘n.s.’: not significant; ‘\**’*: *p* < 0.05; *‘*\*\*’: *p* < 0.01; ‘***’: *p* < 0.001. *N* = 29.

These results indicate that there is significant, widespread information across the brain to allow for the reconstruction of ocular events. Much of what we can see here is interesting as well as intuitive. For example, we see changes in visual activity preceding changes in pupil diameter by 300 ms, despite the absence of external visual events. Blinks are followed 150 ms later by cerebellar-localised responses, and saccades 100 ms later by visual responses. Alongside these clearest signals, we also see a host of spatiotemporal complexity that contributes to the full reconstructable signal.

### Oculomotor Contributions to Auditory Processing

To ask whether oculomotor activity contributes to neural processing in a non-visual cognitive task, we applied the ORFs derived from resting-state data to a separate passive listening paradigm. As a proof-of-principle, all analyses reported in this section were performed at the sensor level. This analysis demonstrates how the ORF method can be used to uncover relationships between oculomotor function and neural activity even in contexts like passive listening, where eye movements are not typically considered. In this task, participants were again instructed to direct their gaze toward a black cross at the centre of a screen while they listened to tone sequences that alternated between predictable (’ordered’) and unpredictable (’random’) patterns.

To investigate the interplay between oculomotor systems and auditory processing, we focused specifically on saccades, motivated by recent findings linking them to excitability in the auditory cortex ^48^. We convolved the saccade time series recorded during the listening task with each participant’s resting-state derived saccade ORFs. This generated a ‘predicted’ time course of saccade-related brain activity. We then applied multivariate pattern analysis (MVPA) to both the actual and this predicted neural activity to uncover the potential contribution of oculomotor dynamics to the processing of auditory regularity (see Fig. 5A).

**Fig. 5:**
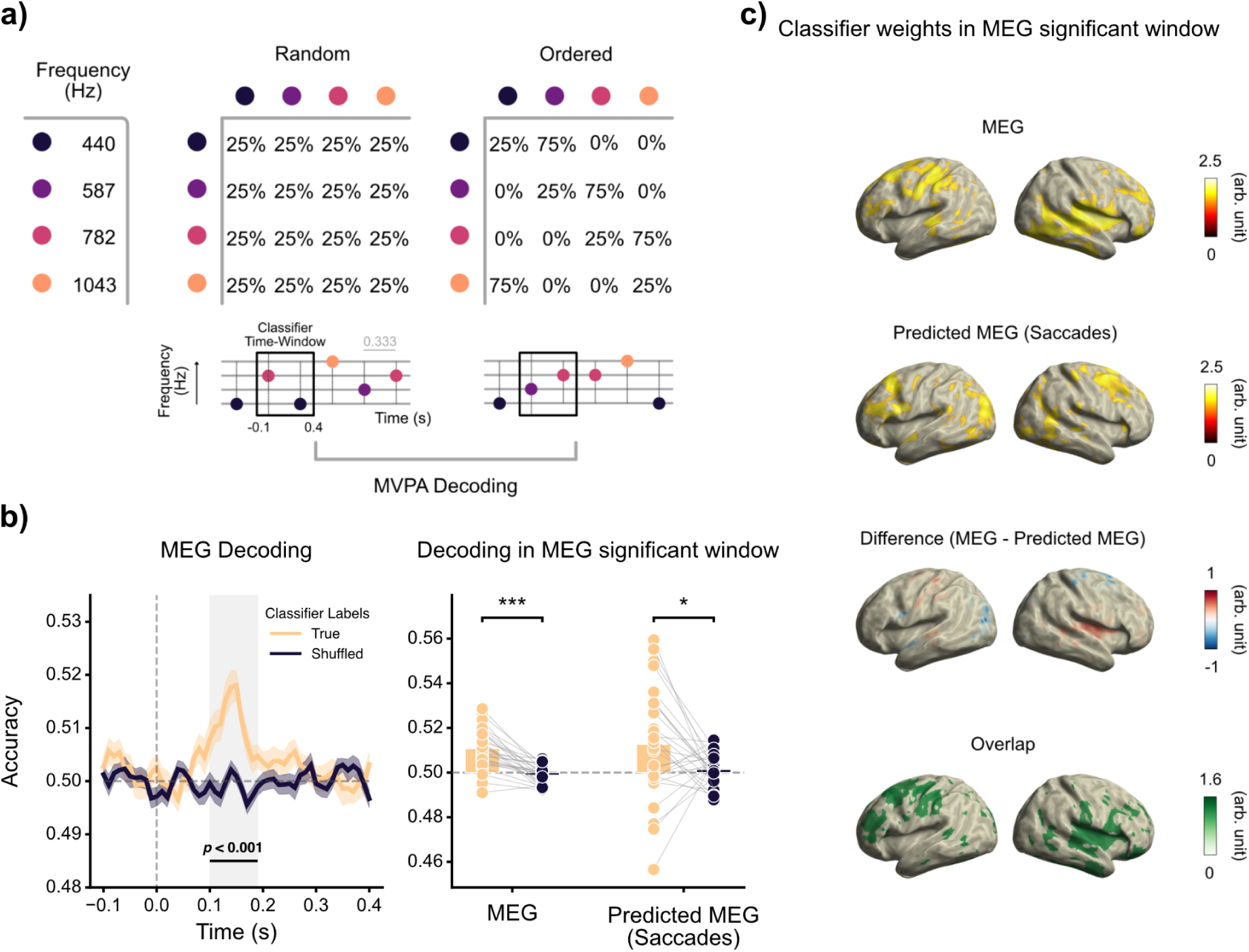
Oculomotor contributions to auditory regularity processing. **a)** Task Schematic. Participants listened to tone sequences that alternated between ordered and random patterns, while instructed to direct their gaze to a central cross. Multivariate pattern analysis (MVPA) was used to decode the regularity condition from the neural data. **b)** MVPA Decoding Performance. Left: time-resolved classification accuracy for the actual MEG (true vs. shuffled labels); the shaded area marks the significant cluster (100–190 ms, p < .001). Right: decoding accuracy averaged within this window for the actual MEG and the predicted saccade-related MEG, each compared against shuffled labels with paired t-tests (MEG: t(27) = 7.13, p < .001, d = 1.69; predicted MEG: t(27) = 2.23, p = .034, d = 0.61). Dots are individual participants. **c)** Source-Projected Classifier Weights. Topographies show brain regions informative for decoding regularity. For actual MEG data, informative patterns originated from the right temporal area and left precentral gyrus. For predicted saccade data, patterns were strongest in the bilateral precentral gyri and left occipital cortex. The overlap highlights shared neural substrates in left frontal and precentral regions, and the right temporal cortex and midbrain. General notes: Weights were averaged across the significant decoding window for MEG (100-190 ms) and thresholded at 75% for visualisation. ‘n.s.’: not significant; ‘\**’*: *p* < 0.05; *‘*\*\*’: *p* < 0.01; ‘***’: *p* < 0.001. *N* = 28.

The MVPA revealed that auditory regularity could be decoded from the actual MEG data and from the predicted saccade-related activity (see Fig. 5B). Cluster-based permutation testing identified above-chance decoding from the actual MEG data between 100 and 190 ms post-tone onset (*p* < .001). Averaging decoding accuracy within this window, classification of true versus shuffled labels was significant for both the actual MEG (*t*(27) = 7.13, *p* < .001, *d* = 1.69) and the predicted saccade-related MEG (*t*(27) = 2.23, *p* = .034, *d* = 0.61) - the latter interestingly reflecting more frequent saccading during the ordered condition (see Supplementary Figures Fig. S4).

To understand the neural basis of this relationship, we projected the MVPA classifier weights into source space for both models and compared their relative spatial distributions (see Fig. 5C). For the actual MEG data, informative activity primarily originated from the right temporal area and the left precentral gyrus. In contrast, the classifier trained on the predicted saccade-related activity drew upon information from the bilateral precentral gyri and the left occipital cortex. A direct comparison highlighted these distinct patterns, with the actual MEG data showing stronger informative activity in the right temporal lobe, while the saccade-predicted data relied more heavily on regions in the right frontal/supplementary motor area and left visual cortex. Crucially, we also observed a substantial overlap between the two patterns in the left frontal and precentral regions, as well as the right temporal cortex and the midbrain. This convergence suggests potential shared neural substrates, where saccade-related dynamics may contribute to the overall neural processing of auditory regularity.

Finally, we assessed how well the time series of each ocular feature could be reconstructed from neural activity recorded during the passive listening task. Note that this reconstruction analysis is distinct from the MVPA classification described above: here, the goal is to predict the continuous or binary time course of an ocular feature, rather than a discrete experimental label. Predictions were generated by convolving the task-related sensor data with the ORFs originally derived from resting-state data. Reconstruction accuracy was then evaluated by correlating the predicted time series with the actual eye-tracking recordings, using Pearson’s *r* for pupil dilation and the Matthews Correlation Coefficient (*MCC*) for blinks and saccades.

As detailed in Figure 6, all ocular features were reconstructed at levels significantly above chance for both the ‘Ordered’ and ‘Random’ conditions. These results demonstrate that the ORFs derived from resting-state data transfer to a different experimental context, capturing oculomotor-related neural dynamics during a cognitive task.

**Fig. 6:**
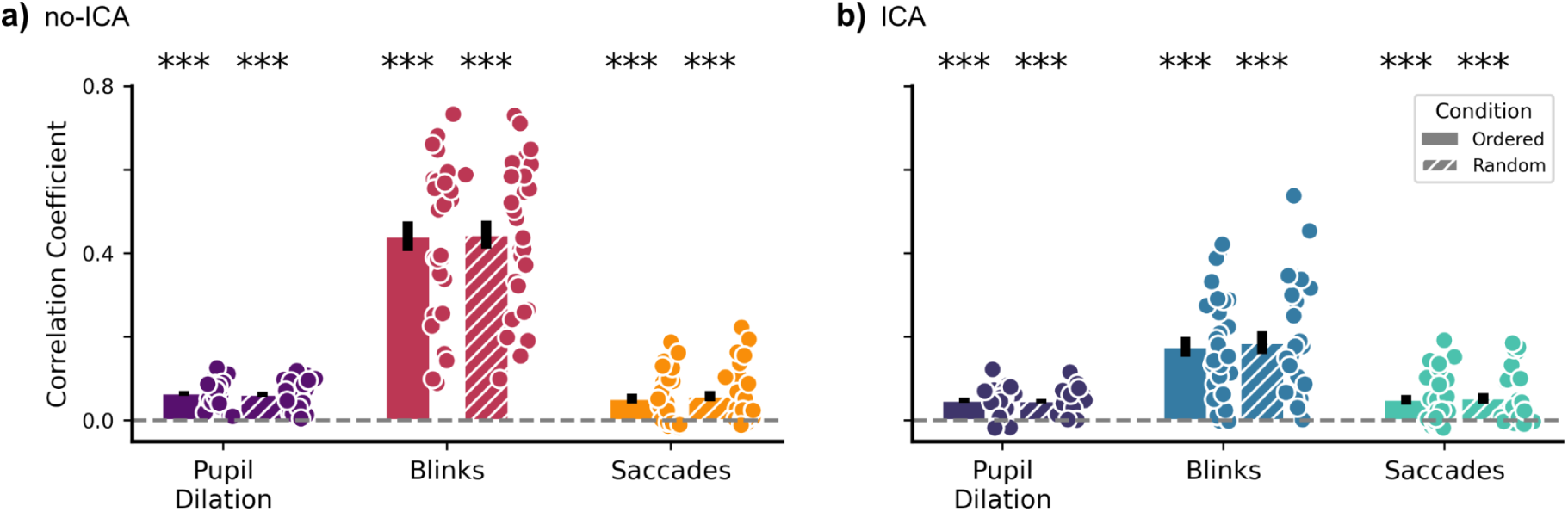
Ocular feature reconstruction from task-related neural activity. Resting-state derived sensor ORFs were used to reconstruct the time series of each ocular feature from neural data recorded during the passive listening task. Performance is shown for the ‘Ordered’ and ‘Random’ auditory conditions. **a)** No-ICA preprocessing: Ordered: Reconstruction was significant for all features (Pupil: *r* = 0.07, *t*(27) = 11.24, *p* < .001, *d* = 2.12; Blinks: *MCC* = 0.44, *t*(27) = 12.47, *p* < .001, *d* = 2.36; Saccades: *MCC* = 0.05, *t*(27) = 4.59, *p* < .001, *d* = 0.87). Random: Reconstruction was also significant for all features (Pupil: *r* = 0.06, *t*(27) = 10.39, *p* < .001, *d* = 1.96; Blinks: *MCC* = 0.44, *t*(27) = 13.17, *p* < .001, *d* = 2.49; Saccades: *MCC* = 0.06, *t*(27) = 4.59, *p* < .001, *d* = 0.87). **b)** ICA preprocessing: Ordered: After ICA, reconstruction remained significant (Pupil: *r* = 0.05, *t*(27) = 8.28, *p* < .001, *d* = 1.56; Blinks: *MCC* = 0.18, *t*(27) = 7.44, *p* < .001, *d* = 1.41; Saccades: *MCC* = 0.05, *t*(27) = 4.32, *p* < .001, *d* = 0.82). Random: Performance was also significant in the random condition (Pupil: *r* = 0.05, *t*(27) = 8.93, *p* < .001, *d* = 1.69; Blinks: *MCC* = 0.19, *t*(27) = 6.83, *p* < .001, *d* = 1.29; Saccades: *MCC* = 0.05, *t*(27) = 4.27, *p* < .001, *d* = 0.81). Statistics were performed using one-sample Student’s t-tests. Error bars reflect standard error of the mean. ‘n.s.’: not significant; ‘\**’*: *p* < 0.05; *‘*\*\*’: *p* < 0.01; ‘***’: *p* < 0.001. *N* = 28.

## Discussion

In this study, we introduce ‘Ocular Response Functions’ (ORF) to characterise the spatiotemporal relationship between neural activity and a comprehensive set of ocular events. By applying a deconvolution approach to resting-state MEG and eye-tracking data, we demonstrated that it is possible to derive anatomically specific transfer functions for saccades, blinks, and pupil dilation. Our results show that these ORFs can predict neural activity from eye movements and, conversely, reconstruct ocular events from brain activity. These analyses revealed several interesting findings. For example, we see changes in visual activity preceding changes in pupil diameter by 300 ms, despite the absence of external visual events, that blinks are followed after 150 ms by cerebellar-localised responses, and saccades after 100 ms by visual responses. Alongside these clearest signals, we also see a range of more subtle factors contributing to the signal with different functional implications. Furthermore, we demonstrated that these functions, derived from task-free data, can be applied to a separate cognitive paradigm to reveal how ocular processes shape task-related neural activity.

Our approach builds upon a well-established tradition of regression-based analyses in neuroimaging ^36–39,49–51^, and more specifically the Temporal Response Function (TRF) framework ^42,43,46^. TRFs are a system identification method typically used to characterise neural responses to complex, external sensory stimuli like natural speech. Our work uses this logic to derive transfer functions from internally-generated oculomotor signals, which we term ‘Ocular Response Functions’ (ORFs). This application also conceptually parallels the use of the hemodynamic response function (HRF) in fMRI analysis, where an impulse response is convolved with an event time series to predict neural activity. However, a key distinction and advantage of our approach is that while the HRF is often assumed to be a canonical, stereotyped function, the ORFs are empirically derived from the data themselves. This allows for a more flexible characterisation of the neural response, revealing distinct transfer functions for different oculomotor events (e.g., blinks vs. saccades).

A further insight concerns the reconstruction of ocular events from neural activity. Recovering sparse, binary events such as saccades and blinks is difficult for standard backward decoding, which is designed for continuous targets and does not cleanly separate correlated features ^47^. Because our reconstruction time-reverses the jointly-estimated forward model, it inherits the feature separation already achieved during encoding, and a subsequent thresholding step - adapted from spike-detection methods in electrophysiology - recovers discrete events directly. Beyond the oculomotor domain, this approach to reconstructing sparse events may be useful wherever response functions are used to model discrete occurrences in continuous neural data, including in speech and language research where such methods were originally developed.

These interpretable transfer functions revealed distinct spatiotemporal profiles for each ocular event, highlighting how the ORF framework can characterise the distributed neural processes underlying oculomotor control. For saccades, the dominant component consistently localised to the occipital visual cortex, peaking at +100 ms - such that, following saccade events, there was a peak in visual processing. This finding confirms the method’s ability to capture neural signatures of visual reafference ^52–56^. For eye blinks, the picture was more multifaceted. After removing the frontocentral ocular artifact via ICA, the dominant ORF component was localised to the cerebellum, while a secondary component revealed activity in visual areas. This source-space dissociation extends EEG studies of the Blink-Related Potential (BRP, e.g., ^57–59)^, which have identified a visual contribution alongside the ocular artefact, by demonstrating the particular contributions also of regions like the cerebellum. This cerebellar localisation warrants caution, of course: deep sources are difficult to resolve with MEG, and signal leakage from other regions cannot be fully excluded. Nonetheless, a cerebellar contribution to blinking is physiologically plausible, given its established role in the timing and execution of eyeblinks ^60,61^ and recent volumetric MEG work indicates that cerebellar sources can be isolated ^62^ . As well as providing better source information via MEG, they also provide an especially flexible tool to complement this classic ERP (or ERF) approach, because they are derived via regression, rather than epoching, approaches. This means that they can describe complex relationships in continuously evolving structures where events (e.g., blinks) can occur in quick succession yet not contaminate overlapping epochs - because there are none.

Similarly, pupil dilation was associated with a distributed network of brain regions, including the visual cortex, cuneus, and temporal lobe. The finding of such widespread, pupil-linked cortical activity using such TRFs, is in line with the comprehensive mapping conducted by Pfeffer et al. ^63^, who used a diverse suite of statistical tools - including cross-correlations and frequency-resolved analyses - to characterise the complex links between pupil-linked arousal and cortical activity. Their work provided a foundational map, detailing the specific lags, frequency-dependencies, and even non-linearities of this relationship within a large resting-state dataset. Our ORF framework advances this important work by first, taking a similar approach to simultaneously isolate responses for blinks and saccades alongside pupillary signals, and second, moving to a unified, predictive model. Rather than employing multiple methods to describe different facets of the relationship, the ORF approach distils these complex dynamics into a single, data-driven linear transfer function, which can act as a reusable toolkit.

One might ask whether the ocular and task contributions could instead be separated within the task, using methods for correlated predictors. The closest is Back-to-Back regression ^47^, which shares the two-step, decorrelating logic of our forward-then-reversed estimation. Such methods adjust each contribution for the other, but face a limit their authors note directly: as factors approach full correlation, no statistical method can separate them and experimental intervention becomes necessary. Stimulus-driven eye movements are especially difficult to therefore study in this way, since the stimulus that drives the cognitive response also triggers the eye movement - thus locking the two in time. Estimating the response function without the stimulus in the present resting state paradigm therefore acts as such an intervention.

One key advantage of this approach is its transferability: the resulting filter can be applied out-of-context to predict oculomotor-related neural activity in new paradigms - a step we illustrate by applying our resting-state ORFs to the passive listening task. The ability of the ORF framework to resolve these anatomically and functionally distinct components provides a potentially useful tool for future studies. It enables researchers to form and test specific hypotheses by interrogating the role of different oculomotor events within targeted brain regions and time windows. More broadly, because the predicted signal is expressed in the units of the neural recording, oculomotor activity can be brought onto the same footing as the brain signal and related to it with the same tools used to study interactions between neural systems - coherence, connectivity, causal modelling, or cross-decoding - opening these analyses to the eye-brain relationship. Deriving this repertoire of distinct, source-localised transfer functions from task-free data provides the foundation for the framework’s key utility: testing how these baseline mappings transfer to new contexts.

The application of our framework to the passive listening task served as a proof-of-principle. The MVPA results, which revealed a spatial overlap between classifier patterns from actual and saccade-predicted neural data, are consistent with a shared neural substrate linking oculomotor control and auditory processing. This was further supported by the reconstruction of all three ocular features from the task data, indicating that the underlying ORF models transfer across different experimental contexts. Ultimately, these findings highlight the framework’s value for uncovering complex, cross-modal contributions of oculomotor systems to cognition. In our test case, frontal, precentral, and temporal regions that could have been believed to be directly related to tone presentation, may in part be shaped by differential oculomotor responses to tone sequences.

While the framework transferred to the listening task, the reconstruction accuracies - especially for pupil dilation and saccades - were modest. This is likely due to two main, deliberate trade-offs in our approach: First, our simple, linear model, chosen for its ability to produce interpretable, spatiotemporally specific transfer functions. While other powerful toolboxes exist for this type of analysis - such as the mTRF-Toolbox ^42,43^ for linear modeling, the unfold toolbox ^40^ for handling non-linearities with generalised additive models (GAMs) or deep learning approaches like cdr(NN) which can capture highly complex, non-linear, and non-stationary dynamics ^64,65^ - we deliberately chose a linear model with a sparsity-promoting boosting algorithm ^41,46^. The choice between different modeling techniques within this family ultimately depends on the scientific goal. If the primary aim is to maximise predictive accuracy - for instance, to reconstruct ocular signals with the highest possible fidelity (e.g., ^66^) - then more complex, non-linear models are likely the superior choice. Our goal here, in contrast, was to prioritise the estimation of interpretable linear transfer functions that can be understood in the same way as neural responses to task events. Second, the ORFs were derived from task-free resting-state data. While this provides an arguably ‘purer’ baseline, the effect sizes would likely be larger during a task. A promising avenue for research is therefore to also compare the ORFs derived here with those estimated during active tasks, such as free viewing or smooth pursuit.

The individual variability we observed in the ORFs may yield informative insights in future work, complementing recent work showing that variability in neurophysiological activity can provide meaningful insights into brain function and personalised neural signatures ^67,68^. In principle, because ORFs yield a quantitative, individual-specific metric, they could be assessed in patient populations and, when combined with existing resting-state datasets, used to compare cohorts. Differences in pupil dilation ORFs between patients with dementia or Parkinson’s disease and healthy controls, for example, might offer insight into altered neural functioning and associated behaviour. Such comparisons would need to account for differences in signal-to-noise ratio and in oculomotor behaviour between groups, but with these caveats, ORFs may offer a basis for developing individualised markers in disorders where oculomotor control is affected.

In summary, we here present Ocular Response Functions (ORF) to characterise and disentangle the distinct neural signatures of different oculomotor events. By deriving interpretable, source-localised transfer functions from task-free resting-state data, our approach provides a baseline for understanding the fundamental mappings between the brain and a repertoire of ocular features. We demonstrated the value of this framework by showing that these baseline ORFs can be applied to probe the contribution of oculomotor dynamics during a separate cognitive task. This work underscores the importance of modeling, rather than simply removing, ocular-related signals and provides an accessible framework to investigate their intricate interplay with neural processing in both cognitive and clinical neuroscience.

## Methods

### Participants

We used data from 29 participants (12 female, 17 male; mean age = 25.70, range = 19 - 43). For the passive listening task, the eye tracker failed to detect the pupil and corneal reflection for one participant, leading to a flat signal for the majority of one block. As a result, no valid data were recorded for that block and the participant was excluded from the analysis, leaving a total of 28 participants for the passive listening task (29 remain in the core reported analyses). All participants reported normal hearing and had normal, or corrected to normal, vision. They gave written, informed consent and reported that they had no previous neurological or psychiatric disorders. The experimental procedure was approved by the ethics committee of the University of Salzburg and was carried out in accordance with the declaration of Helsinki. All participants received either a reimbursement of 10 € per hour or course credits.

### Stimuli and Experimental Design

We assessed participants’ head shapes before the start of the experiment using cardinal head points (nasion and pre-auricular points), digitised with a Polhemus Fastrak Digitiser (Polhemus), and around 300 points on the scalp. For all parts of the experiment, for both the resting-state period and the passive tone listening task, participants were instructed to direct their gaze toward a black cross at the centre of a screen. Each MEG session then started with a 5-minute resting-state recording. This standard resting-state paradigm proved ideal for our goal of establishing fundamental ORFs in the absence of complex visual inputs or cognitive demands, which could otherwise confound the eye-brain relationship. While this task minimises large, exploratory saccades, it provides a clean baseline and generates ample data on blinks, microsaccades, and pupil fluctuations for our models. During the 5-minute resting-state period, participants produced an average of 0.32 ± 0.24 Hz for blinks and 1.80 ± 1.26 Hz for saccades. Afterwards, the individual hearing threshold was determined using a pure tone of 1043 Hz. This was followed by 2 blocks of passive listening to tone sequences of varying entropy levels. Sequences consisted of four different pure tones (f1: 440 Hz, f2: 587 Hz, f3: 782 Hz, f4: 1043 Hz, each lasting 100 ms), each block consisted of 1500 tones presented with a temporally predictable rate of 3 Hz. Entropy levels (ordered / random) changed pseudorandomly every 500 trials within each block, thus resulting in a total of 1500 trials per entropy condition. While in an ‘ordered’ context forward transitions (i.e. f1→f2, f2→f3, f3→f4, f4→f1) had a high probability of 75%, repetitions (e.g., f1→f1, f2→f2,…) were rather unlikely with a probability of 25%. However, in a ‘random’ context all possible transitions (including forward transitions and repetitions) were equally likely with a probability of 25%. Oculomotor behavior during this passive listening task was largely comparable to the resting state, with average rates of 0.30 ± 0.22 Hz for blinks and 1.59 ± 0.87 Hz for saccades.

### Data Acquisition and Preprocessing

#### MEG

We collected MEG data using a whole head system (Elekta Neuromag Triux, Elekta Oy, Finland), placed within a standard passive magnetically shielded room (AK3b, Vacuumschmelze, Germany). The signal was recorded with 102 magnetometers and 204 orthogonally placed planar gradiometers at 102 different positions at a sampling frequency of 1 kHz (hardware filters: 0.1 - 330 Hz). In a first step, a signal space separation algorithm, implemented in the Maxfilter program (version 2.2.15) provided by the MEG manufacturer, was used to clean the data from external noise and realign data from different blocks to a common standard head position. Data preprocessing was performed using Matlab R2020b (The MathWorks, Natick, Massachusetts, USA) and the FieldTrip Toolbox (Oostenveld et al., 2011). All data were band-pass filtered between 0.5 Hz and 30 Hz (onepass-zerophase finite impulse response (FIR) filter, order 3330, hamming-windowed) and downsampled to 100 Hz. We further used two versions of preprocessed data that either used independent component analysis to reject eye movement (and heart-rate) related artefacts or not. We further refer to these data as either ICA or no-ICA respectively. For ICA, to identify eye movement and heart rate artefacts, 65 independent components were identified from filtered (onepass-zerophase lowpass FIR filter, order 132, hamming-windowed; onepass-zerophase highpass FIR filter, order 1650, hamming-windowed) continuous data of the resting-state recordings using runica-ICA. The goal of this ICA step was to remove non-neurogenic artifacts while preserving the underlying neural signals. Based on visual inspection of component time-courses and spatial topographies, we identified and removed components dominated by cardiac or by ocular activity, including that from the eyelids (blinks), eyeball rotation, and extraocular muscles. This process resulted in the removal of a median of 3 components per participant (min = 2, max = 4).

#### Eye Tracking

We used a Trackpixx3 binocular tracking system (Vpixx Technologies, Canada) with a 50 mm lens. We synchronised MEG and eye-tracking data feeding the Trackpixx’s output data as three additional channels into the MEG system, representative of horizontal gaze, vertical gaze, and pupil dilation of participants’ right eye. Recordings started after a 13-point calibration and validation procedure. Throughout the experiment participants were seated in the MEG at a distance of 82 cm from the screen, with their chin resting on a chinrest to reduce head movements. We used an algorithm based on pupillometry noise to detect blinks ^69^. To account for potential residual blink artefacts when examining pupil dilation, we removed an additional 100 ms around blink edges. Blinks were then removed from pupil dilation data and linearly interpolated. For saccade detection, we used a velocity-based detection approach ^2–4^ using the Euclidean distance between successive gaze points in the horizontal and vertical planes as in ^5^ with a 7 ms smoothing window on the resulting velocity vector (using the built “smoothdata” function in Matlab), thus considering two-dimensional velocity profiles. Saccade on-/offsets (i.e. durations) were then identified where velocity exceeded and subsequently fell below the threshold of five times the median velocity. A minimum delay of 100 ms was used between saccade onsets to prevent multiple detections of the same saccade. We additionally excluded saccades from pupil dilation data and linearly interpolated the gaps. For further analysis, blinks and saccades were transformed into binary vectors of zeros (no blink/saccade) and ones (blink/saccade) retaining event durations. Afterwards, pupil dilation data were bandpass filtered between 0.01 - 4 Hz (onepass-zerophase finite impulse response (FIR) filter, order 165000, hamming-windowed) and median centred on the screen centre. Finally, the three eye tracking data channels (pupil dilation, blinks, and saccades) were appended to the MEG data and therefore downsampled to 100 Hz alongside the MEG data. Note that binary blink and saccade vectors were then restored by adding “1 s” at the nearest time points of the original sampling rate to avoid potential data loss of short activation patterns (i.e. onsets).

#### MEG and Eye Tracking

Resting-state data were cut from 10 s to 290 s after recording onset. Data of the entropy modulation paradigm were cut from −1 s of the first tone to +1 s of the last tone of every 500 tone segment and corrected for a 16 ms delay between trigger and stimulus onset generated by sound travelling through the pneumatic headphones into the shielded MEG room.

### The Ocular Response Function

#### Encoding Models

To establish spatially-resolved transfer functions from oculomotor action to neural representations, we first projected the continuous resting-state MEG data into anatomical source space (see Source Projection). We then estimated a multivariate temporal response function (mTRF) for the time series of each source-space voxel using the Eelbrain toolkit ^41^. A deconvolution algorithm based on boosting ^46^ was applied to the source-space data to estimate the optimal TRFs that predict neural activity (dependent variable) from three key ocular features (predictors): pupil dilation, blinks, and saccades. We deliberately chose boosting over ridge regression ^42,43^ as it imposes a sparse prior, encouraging TRFs with fewer non-zero elements. This sparsity helps in interpreting the core temporal components of the response. For model fitting, the predictor time series and the neural time series for each voxel were separately centered and standardised. The model was trained to find the optimal linear filter over time lags from −1 s to +1 s, using a 100 ms wide Hamming window basis (please refer to ^41^ for further details on the boosting algorithm and parameters). A 4-fold cross-validation approach was used to evaluate the model and avoid overfitting ^70^. All predictors were included into the same encoding model. All three predictors were included in the same encoding model, and we used selective stopping to prevent overfitting on individual predictors while allowing others to continue training ^41^. This process results in a unique ORF for each ocular feature at each source-space voxel.

#### Reconstruction and Evaluation Metrics

For reconstruction, the estimated ORF filters are simply reversed in time to predict ocular features from brain activity. This time-reversal is a computationally efficient method, analogous to a cross-correlation, for assessing the bidirectional relationship using the same estimated kernel.

Model performance was evaluated on held-out test data from the cross-validation procedure. For reconstruction, since blinks and saccades are binary events, the continuous model predictions were converted into probabilities using a sigmoid function and then thresholded at 0.5. We then calculated the Matthews Correlation Coefficient (MCC) to assess performance for blinks and saccades ^71,72^ and Pearson’s correlation (r) for the continuous pupil dilation signal. The same correlation metrics were used to evaluate encoding performance.

### Source projection

To perform our main analyses in anatomical source space, we projected the continuous MEG time series data using Linearly Constrained Minimum Variance (LCMV) spatial filters ^73^. LCVM filters were computed by warping anatomical template images to the individual head shape and further brought into a common space by co-registering them based on the 3 anatomical landmarks (nasion, left, and right preauricular points) with a standard brain from the Montreal Neurological Institute (MNI, Montreal, Canada; ^74^). Then, we computed a single-shell head model ^75^ for each participant. As a source model, a grid with 1 cm resolution and 2982 voxels based on an MNI template brain was morphed into the brain volume of each participant. This allowed group-level averaging and statistical analysis as all the grid points in the warped grid belong to the same brain region across subjects. Spatial filters were estimated by following a publicly available tutorial that was made available by FieldTrip and can be found on their homepage (https://www.fieldtriptoolbox.org/workshop/paris2019/handson_sourceanalysis/). A key step in our procedure was to utilise separate empty-room recordings to compute a robust noise covariance matrix. The continuous resting-state MEG data were then prewhitened using this noise covariance to suppress environmental and sensor noise ^76^. The dimensionality reduction parameter for this step (kappa) was determined automatically by detecting the first major “cliff” in the singular value spectrum of the noise covariance matrix.

Following prewhitening, a data covariance matrix was computed from the main experimental data. A forward model (leadfield) was then prepared using the subject-specific headmodel and source model grid. Finally, the LCMV spatial filters were computed with a regularisation parameter (lambda) set to 5%. We specified a fixed dipole orientation (fixedori = ‘yes’) and used a ‘unit noise gain’ normalisation (weightnorm = ‘unitnoisegain’) to ensure uniform noise projection across all source locations. This entire procedure was performed separately for data with and without prior ICA correction, yielding two sets of spatial filters for each participant. These filters were then used to project the continuous resting-state MEG data into source space for subsequent ORF analyses. Anatomical labels for source-space results were obtained using the AAL3 ^77^. Wherever the coordinates of an effect in mni-space did not correspond to any labelled region, nearest neighbour interpolation was applied. This approach allowed for the assignment of unlabeled coordinates by identifying the closest labelled region.

### Principal component analysis

To reduce the dimensionality of the data and identify the most relevant spatiotemporal components of the neural response, we applied principal component analysis (PCA). This was performed on the source-projected ORFs for each subject and ocular feature individually, transforming the high-dimensional voxel space into a smaller set of orthogonal components that capture the most variance in the eye-brain relationship.

The analysis pipeline was as follows: First, to mitigate potential edge artifacts from the deconvolution, we cropped the ORFs to a time window of −0.9 to +0.9 seconds. To focus the decomposition on response magnitude irrespective of direction, we then computed the absolute value of these cropped ORFs. PCA was then performed on this rectified (absolute-magnitude) data. Following the decomposition, a polarity alignment procedure was applied to ensure component consistency across subjects. Because the sign of a principal component is arbitrary, we standardised the polarity by projecting each component’s spatial weights back onto the original magnitude data. If this projection resulted in a negative sum, the signs of both the spatial weights and the temporal component were inverted. This step ensures that the directionality of the main effect (e.g., a positive peak) is consistent across all participants before averaging or group-level analysis. For all subsequent analyses, we consistently selected the top three principal components that explained the most variance for each ocular feature and preprocessing condition (ICA vs. no-ICA).

### Multivariate Pattern Analysis (MVPA)

To gain deeper insights into how oculomotor activity may influence neural activation patterns during experimental tasks, we applied sensor-level resting-state derived ORFs to data from a passive listening paradigm (see Fig. 5A; ^7^). Given recent findings linking neural excitability in the auditory cortex with saccades ^48^, we focused on potential overlaps with saccade-related neural activity. For reasons of computational efficiency and because this constituted solely a proof-of-principle, this specific analysis was performed entirely on sensor-level data. We estimated sensor-space saccade ORFs from the resting-state recordings and then convolved them with the saccadic eye movement data from the passive listening task, generating a predicted MEG time series for all 306 sensors.

Both the ICA-cleaned MEG and predicted MEG data were epoched around tone onsets (−0.1 to 0.4 seconds), resulting in 3000 epochs (1500 for the ordered regularity condition and 1500 for the random condition). The data were subjected to time-resolved classification using the MVPA-Light package ^78^. We employed the default 5-fold cross-validation, z-scoring as a preprocessing step, and linear discriminant analysis (LDA) to decode regularity conditions (i.e. ordered vs. random) from both MEG and predicted MEG data, as well as a shuffled version of those labels as a control. This process yielded one accuracy value per time point and one classifier weight per time point per sensor for each participant.

Classifier weights were corrected by their covariance structure ^79^, projected into source space by multiplying them with the resting-state spatial filters, and baseline-corrected (relative change) from −0.05 - 0 s ^80^ relative to tone onset. We then averaged these weights from 0.10 - 0.19 s (sig. Cluster of the MEG decoding, see Fig. 5B). To ensure comparability between true and predicted MEG results, whilst maintaining the relative importance of brain areas in sensitivity to changes in stimulus regularity, weights from both datasets were z-scored (i.e., standardised to a mean of 0 and standard deviation of 1). After this transformation, the negative of the minimum value across both datasets (i.e. global minimum) was added to retain zero values (z’ = z + (min(z) * −1)).

To explore the shared contributions of neural and oculomotor processes to regularity processing, we calculated both the difference and overlap of the averaged, normalised classifier weights. The difference was computed by subtracting the predicted MEG weights (Weights_PredictedMEG_) from the MEG weights (Weights_MEG_). The overlap was computed according to the following formula:

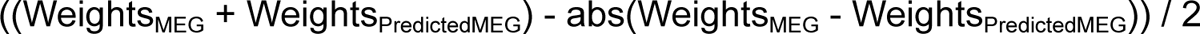

### Statistical Analysis

For the statistical analysis of resting-state data, we employed one-sample Student’s t-tests to evaluate the significance of the encoding and reconstruction correlation coefficients (see evaluation metrics). To obtain one encoding correlation coefficient for each participant, we first averaged the PCA weight matrices across participants and identified the voxel with the maximum component weight for each of the first three components. We then extracted the correlation coefficient from these peak locations for each participant. Subsequently, we report the r-values, t-values, p-values, and Cohen’s d effect sizes for each one-sample Student’s t-test. We intentionally chose to use one-sample t-tests over alternatives such as ANOVA or mixed-effects models, as we did not aim to assess whether one type of eye movement was more strongly represented in neural activity than another. Our primary interest was to demonstrate that ocular behaviour is consistently represented across various brain regions, allowing for reliable encoding and reconstruction effects, rather than exploring the extent of differences between them.

To statistically evaluate the MVPA analysis of the passive listening data, we used a cluster-based randomisation approach. This analysis was conducted across all time points within the classifier time window (−0.1 to 0.4 seconds), comparing classifier accuracy for true versus shuffled regularity condition labels (ordered vs. random) for the actual MEG data (see Fig. 5B). We computed the randomisation distribution of t-values across 10,000 permutations, using a cluster-forming alpha threshold of 0.05. The resulting clusters were compared against the original contrast at an alpha level of 0.05. For each significant cluster, we report the average t-values, p-values, and Cohen’s d (effect size) across the time points within the cluster. To then compare the actual and predicted saccade-related MEG on the same footing, we averaged each participant’s classification accuracy across this significant cluster window and compared true versus shuffled labels using paired-samples t-tests, computed separately for the actual MEG and the predicted saccade-related MEG (reporting t, degrees of freedom, p, and Cohen’s d). Decoded eye movements and the corresponding correlation coefficients were evaluated in the same manner as for the resting-state data.

## Data availability

Preprocessed Data required to reproduce the analyses supporting this work are publicly available in the corresponding author’s Open Science Framework repository (https://osf.io/5bucm/). Raw data (> 250 GB) will be shared upon request.

## Code availability

Code to analyse preprocessed data and further reproduce results and figures from this manuscript is available in the corresponding author’s Open Science Framework repository (https://osf.io/5bucm/).

## Acknowledgements

Thanks to the whole research team. Special thanks to Manfred Seifter for his support in conducting the measurements, to Gareth Barnes for comments on the manuscript, and to the MEG group at the Functional Imaging Lab, UCL, for input on the idea. This work was supported by a Leverhulme Project Grant (RPG-2022-358) and European Research Council (ERC) consolidator grant (101001592) under the European Union’s Horizon 2020 research and innovation programme, both awarded to C.P.

## Author contributions

Q.G. designed the experiment, collected and analysed the data, generated the figures, and wrote the manuscript. J.S. designed the experiment, collected and analysed the data, and edited the manuscript. A.K. contributed to data analyses and edited the manuscript. N.W. collected the data, contributed to data analyses and edited the manuscript. C.P. acquired the funding, supervised the project, and edited the manuscript.

## Notes

### Competing Interest Statement

The authors have declared no competing interest.

### Summary of Updates

Clarification on approach vs method in general; Figure 5b updated to show MVPA decoding within significant MEG decoding

