## Supplemental information for "Ocular Response Functions reveal how ocular processes relate to neural activity"

### 1 Supplementary Material

#### 2 Supplementary Figures

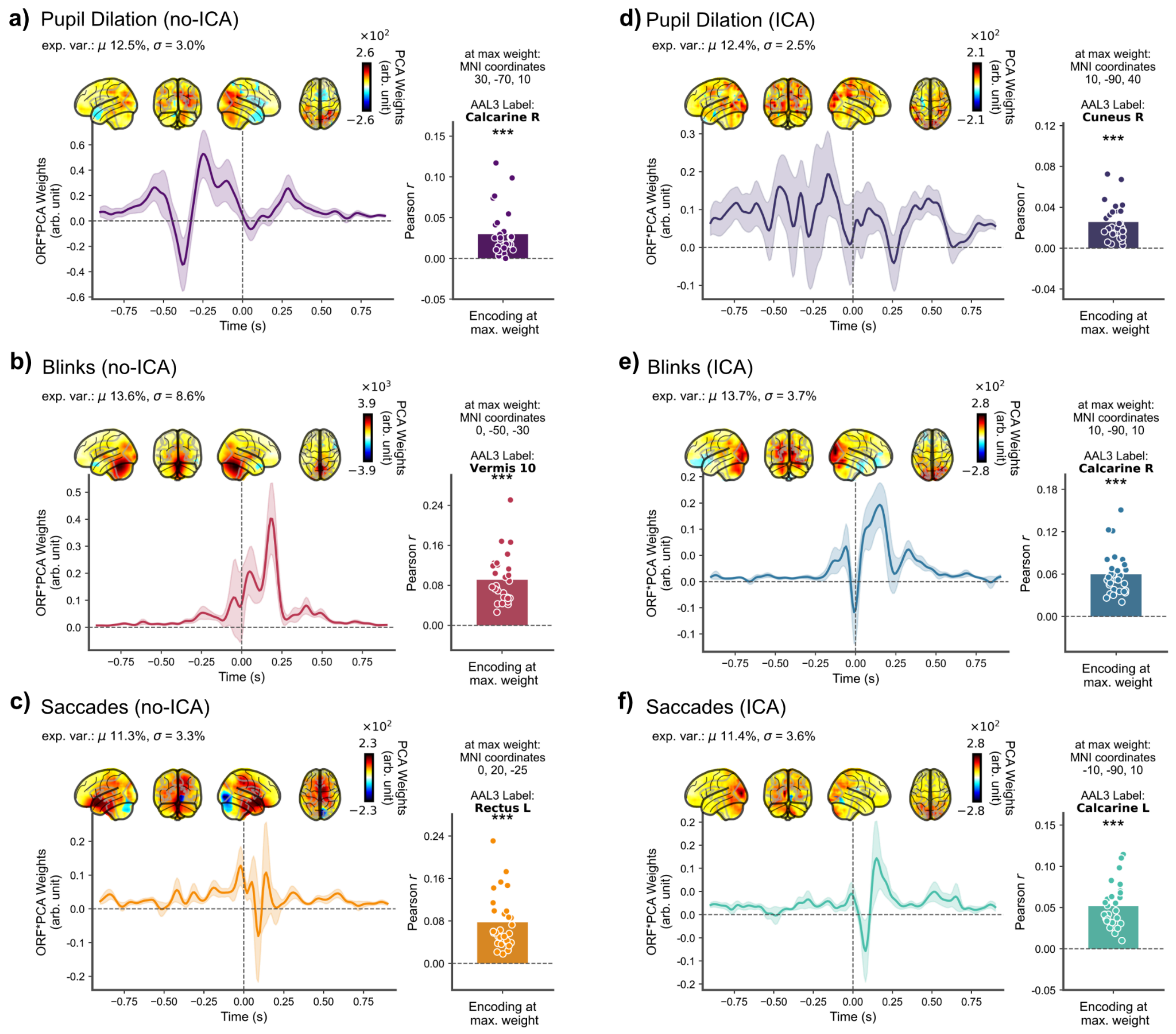


**Fig. S1:** *Encoding results for the second principal component (PC2) of the Ocular Response Functions*. Results are shown for data without (no-ICA) and with (ICA) oculomuscular artifact correction. **a-c)** No-ICA Results: **a)** Pupil dilation: The component showed primary loadings over visual areas and peaked at ~ -250 ms. **b)** Eye blinks: The component was localised to cerebellar regions and peaked around +200 ms. **c)** Saccades: The component loaded maximally over frontal regions with a more distributed spatial and a more complex temporal pattern. A small peak can be related to frontal loadings approximately at 0 ms. **d-f)** ICA-Cleaned Results: **d)** Pupil dilation: After artifact correction, the component's primary loading shifted to occipital visual areas with additional contributions from distributed sources, leading to no clear temporal peak but instead a subset of peaks representative of those distributed networks. **e)** Eye blinks: The component was now localised to the visual cortex and exhibited a peak around +200 ms. **f)** Saccades: The component remained localised to the occipital visual cortex with a peak at ~ +100 ms, confirming a robust, non-muscular neural source. *General Notes*: Shaded error bars represent 95% CI's. Anatomical labels were obtained using the automated anatomical labelling atlas 3 (AAL3). Statistics were performed using one-sample Student's t-tests. 'n.s.': not significant; '*': $p < 0.05$; '**': $p < 0.01$; '***': $p < 0.001$. $N = 29$.

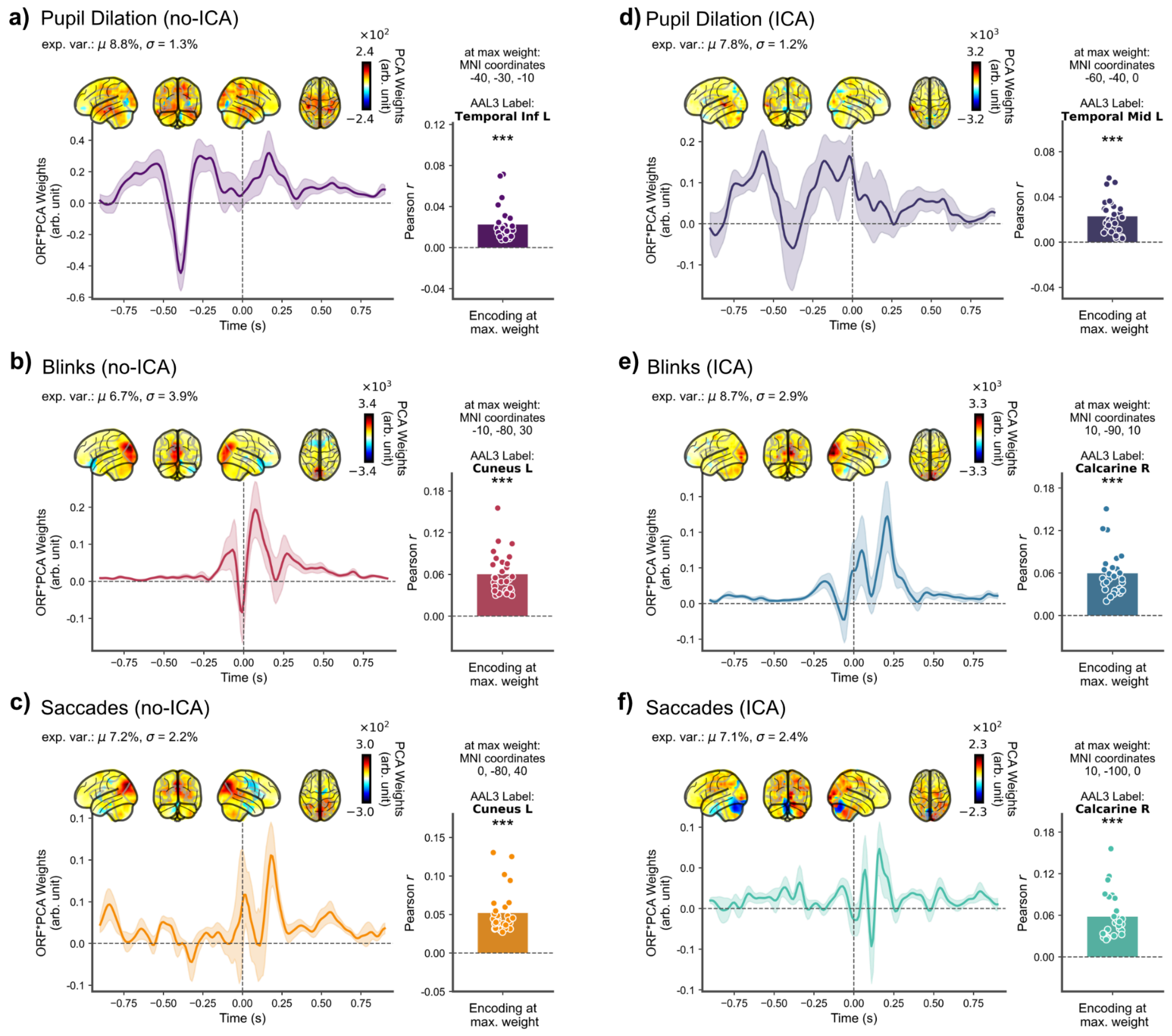


**Fig. S2:** *Encoding results for the third principal component (PC3) of the Ocular Response Functions*. Results are shown for data without (no-ICA) and with (ICA) oculomuscular artifact correction. **a-c)** No-ICA Results: **a)** Pupil dilation: The component showed primary loadings over temporal areas and exhibited a negative peak at ~ -400 ms. **b)** Eye blinks: The component was localised to visual regions and peaked around +100 ms. **c)** Saccades: The component loaded maximally over visual regions with a more complex temporal pattern. The main peak can be observed at approximately +200 ms. **d-f)** ICA-Cleaned Results: **d)** Pupil dilation: After artifact correction, the component's primary loading remained over (more left lateralised) temporal areas. A dual peak pattern can be observed at about -550 and -200-0 ms. **e)** Eye blinks: The component was again localised to the visual cortex and exhibited a peak around 200 ms. **f)** Saccades: The component remained localised to the occipital visual cortex with a less pronounced peak at ~ +200 ms, confirming a robust, non-muscular neural source. *General Notes*: Shaded error bars represent 95% CI's. Anatomical labels were obtained using the automated anatomical labelling atlas 3 (AAL3). Statistics were performed using one-sample Student's t-tests. 'n.s.': not significant; '*': $p < 0.05$; '**': $p < 0.01$; '***': $p < 0.001$. $N = 29$.

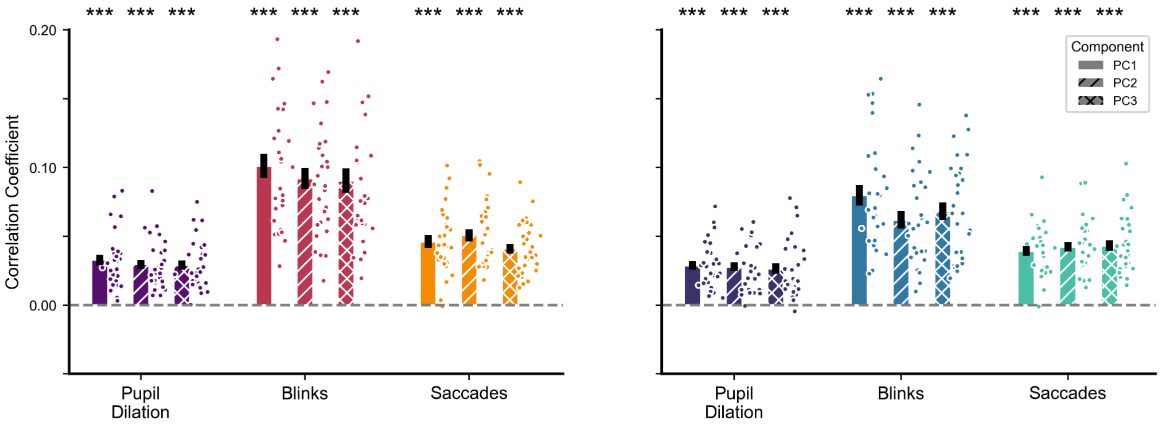


**Fig. S3:** *Reconstruction performance for principal components of ocular features from resting-state data.* Reconstruction was performed using source-space ORFs weighted by the first (PC1), second (PC2), or third (PC3) principal component. **a)** No-ICA preprocessing: All three principal components yielded significant decoding for all features. PC1: Pupil: $r$ = 0.03, $t$(28) = 8.66, $p$ < .001, $d$ = 1.61; Blinks: $MCC$ = 0.10, $t$(28) = 11.62, $p$ < .001, $d$ = 2.16; Saccades: $MCC$ = 0.05, $t$(28) = 10.25, $p$ < .001, $d$ = 1.90. PC2: Pupil: $r$ = 0.03, $t$(28) = 8.89, $p$ < .001, $d$ = 1.65; Blinks: $MCC$ = 0.09, $t$(28) = 11.94, $p$ < .001, $d$ = 2.22; Saccades: $MCC$ = 0.05, $t$(28) = 11.25, $p$ < .001, $d$ = 2.09. PC3: Pupil: $r$ = 0.03, $t$(28) = 8.65, $p$ < .001, $d$ = 1.61; Blinks: $MCC$ = 0.09, $t$(28) = 10.12, $p$ < .001, $d$ = 1.88; Saccades: $MCC$ = 0.04, $t$(28) = 11.79, $p$ < .001, $d$ = 2.19. **b)** ICA preprocessing: After ICA, decoding performance for all components remained highly significant. PC1: Pupil: $r$ = 0.03, $t$(28) = 9.31, $p$ < .001, $d$ = 1.73; Blinks: $MCC$ = 0.08, $t$(28) = 10.69, $p$ < .001, $d$ = 1.99; Saccades: $MCC$ = 0.04, $t$(28) = 11.32, $p$ < .001, $d$ = 2.10. PC2: Pupil: $r$ = 0.03, $t$(28) = 8.66, $p$ < .001, $d$ = 1.61; Blinks: $MCC$ = 0.06, $t$(28) = 9.62, $p$ < .001, $d$ = 1.79; Saccades: $MCC$ = 0.04, $t$(28) = 12.21, $p$ < .001, $d$ = 2.27. PC3: Pupil: $r$ = 0.03, $t$(28) = 6.88, $p$ < .001, $d$ = 1.28; Blinks: $MCC$ = 0.07, $t$(28) = 10.37, $p$ < .001, $d$ = 1.93; Saccades: $MCC$ = 0.04, $t$(28) = 11.01, $p$ < .001, $d$ = 2.05). Statistics reflect one-sample Student's t-tests against zero. Error bars reflect standard error of the mean. 'n.s.': not significant; '*': $p$ < 0.05; '**': $p$ < 0.01; '***': $p$ < 0.001. $N$ = 29.

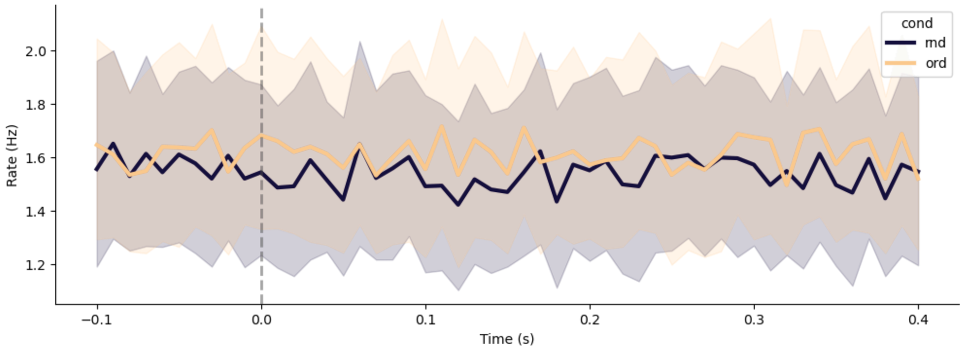


**Fig. S4:** *Saccade rates across participants for the two regularity conditions of the passive listening task.* In general, participants appeared to engage slightly more in saccadic eye movements during the ordered (ord) condition (yellow) compared to the random (rnd) condition (purple). However, this pattern was accompanied by considerable variability across participants and over time. Shaded error bars represent 95% CI's. $N$ = 28.
